# Cryogenic electron tomography and elemental analysis of mitochondrial granules in human retinal ganglion cells

**DOI:** 10.1101/2024.12.17.628994

**Authors:** Gong-Her Wu, Cathy Hou, Andrew Thron, Hirenkumar Rajendra Patel, Liam Spillane, Sanket Rajan Gupte, Serena Yeung-Levy, Sahil Gulati, Christopher Booth, Yaping Joyce Liao, Wah Chiu

## Abstract

Combining three-dimensional (3D) visualization with elemental analysis of vitrified cells can provide crucial insights into subcellular structures and elemental compositions in their native environments. We present a coordinated approach using cryogenic electron energy loss spectroscopy (cryoEELS) and cryogenic electron tomography (cryoET) to characterize the elemental distribution and ultrastructure of vitrified cells. We applied this method to examine calcium disposition in the mitochondria of cultured human retinal ganglion cells (RGCs) exposed to pro-calcifying conditions relevant to optic disc drusen pathology. Our cryoEELS analysis revealed mitochondrial granules with elevated calcium signals, offering direct evidence of mitochondrial calcification. Additionally, cryoET coupled with artificial intelligence-based analysis enabled quantification of the volume and spatial distribution of these calcium granules. This integrated workflow can be broadly applied to various cell types, facilitating the study of ultrastructure and elemental distribution in subcellular structures under diverse physiological and pathological conditions, as well as in response to therapeutic interventions.

## Introduction

Calcium ions play a critical role in various cellular functions, including signal transduction, energy production, and cell death regulation ^1^. Mitochondria are essential in buffering and modulating cellular calcium levels by controlling the uptake and release of these ions ^2,3^. While the intake of mitochondrial calcium stimulates oxidative metabolism and ATP synthesis, enhancing cellular energy production, excessive calcium accumulation within mitochondria can trigger apoptotic cell death ^4^. Recent advances in molecular biology have identified the complex machinery involved in mitochondrial calcium transport ^5–7^ and have linked disrupted mitochondrial calcium homeostasis to various diseases, such as neurodegenerative disorders ^8^ and cardiac dysfunction ^9^. Understanding the molecular mechanisms underlying mitochondrial calcium regulation is crucial for developing targeted therapeutic strategies.

Using electron microscopy, mammalian mitochondria in chemically fixed cells have been found to accumulate calcium phosphate precipitates ^10,11^. Additionally, the composition of dense granules isolated from mitochondria were determined to be calcium and phosphate ^12^. Wolf et al. used cryogenic scanning transmission electron tomography (STEM) and cryogenic energy-dispersive X-ray (EDX) spectroscopy to demonstrate the presence of calcium and phosphate within regions of the mitochondrial matrix containing electron-dense granules in vitrified samples ^13^.

Electron energy loss spectroscopy (EELS) is known to offer advantages over EDX for characterizing the elemental composition of specimens at the nanometer scale when incorporated with traditional transmission electron microscopy (TEM) or STEM ^14^. EELS is a more dose-efficient technique compared to EDX for spectral features <2000 eV since a higher number of forwarded scattered electrons are collected in the post-column spectrometer compared to the percentage of collected X-rays that are emitted from the sample. This provides a higher signal-to-noise ratio in the collected EELS spectra than the EDX spectra for an equal acquisition time. Due to the higher energy resolution (<1 eV) of EELS, compared to EDX, EELS exhibits increased sensitivity to lighter elements, especially carbon, nitrogen, and oxygen. This is also true for elements that have prominent L_2,3_ edges (white lines) such as calcium and iron ^15^. While both techniques can achieve sub-nanometer spatial resolution ^16,17^, EDX requires a longer acquisition time resulting in higher cumulative doses, during which a sample is more likely to be damaged. EELS, on the other hand, delivers finer spatial resolution at a lower total dose ^18,19^, making it a powerful tool for the compositional analysis of dose-sensitive materials.

Due to the low detective quantum efficiency (DQE) of photodiode array detectors, elemental mapping in biological samples was more dose-efficient with energy-filtered transmission electron microscopy (EFTEM) ^20,21^. Furthermore, the significant increase in DQE with the implementation of the charge-coupled device (CCD) camera could decrease the total dose needed to acquire STEM-EELS spectrum images (SI) by an order of magnitude compared to EFTEM imaging ^22,23^. STEM-EELS SI has been successfully used to characterize chemical segregation of trace elements in chemically fixed and dehydrated tissue samples, as well as isolated proteins and macromolecules^24–26^. However, the electron dose used to collect the data for the core-loss edges (>100 eV) is typically greater than ≥ 10^7^ e^-^/Å^2^ which would damage vitrified samples. For instance, above 100 e^-^/Å^2^, the formation of H_2_ bubbles was observed. Such dose has limited the analysis to using valence EELS (<100 eV) for vitrified samples ^27^. This greatly hinders the elemental analysis due to the limited accessible spectral range and the strongly excited bulk plasmon which overlaps with the valence edges.

The introduction of radiation-hardened, direct detection CMOS cameras to single particle imaging enabled researchers to push the spatial resolution of particle reconstruction due to a >2x increase in DQE ^28^. When applying direct detection such as K3 camera to EELS, the same increase in DQE results in an almost 3-fold increase in the signal-to-noise ratio observed for weak, core-loss edges ^29^. The increase in sensitivity has already been shown to double the spatial resolution of EELS SI acquired from synthetically derived polymers, while reducing the dose needed to 10 e^-^/Å^2 30^. Such a dose-efficient detector opens the possibility of applying cryogenic electron energy loss spectroscopy (cryoEELS) to vitrified biological specimens.

Here, we demonstrate the quantitative abilities of cryoEELS and cryogenic electron tomography (cryoET) for direct elemental analysis and 3D visualization of subcellular structures, respectively, in vitrified biological samples. Specifically, we apply our workflow to study mitochondrial calcium accumulation in human retinal ganglion cells (RGCs) under pro-calcifying conditions. RGCs have been implicated in optic disc drusen (ODD), the most common cause of optic nerve ectopic calcification, leading to vision loss ^31^. Clinically, ODD is characterized by refractile, heavily calcified extracellular concretions of various sizes in the unmyelinated anterior optic nerve–a location with a high concentration of mitochondria ^32^. Ultrastructurally, optic nerves in ODD autopsy eyes contain intra- and extra-axonal calcified mitochondria ^33^. These calcified deposits consist mainly of calcium phosphate with trace amounts of amino acids, nucleic acids, mucopolysaccharides, glycoproteins, and mitochondrial fragments ^34^. While the underlying mechanisms of ODD remain unknown, studies have proposed that impaired axonal metabolism, which leads to axonal degeneration and extrusion of calcified mitochondria into the extracellular space, may contribute to its pathophysiology ^33,35,36^.

To develop an *in vitro* cellular model of ODD, we cultured human embryonic stem cell-derived RGCs treated with phosphates–a well-established model for inducing ectopic calcification based on the leading theory for pathogenesis of atherosclerosis and other diseases ^37^. Phosphate was supplied in two different ways: inorganic phosphate in the form of potassium phosphate monobasic, KH_2_PO_4_, (PP) and organic phosphate in the form of a cocktail of β-glycerophosphate, dexamethasone, ascorbic acid, hereafter known as calcification media (CM). Inorganic phosphate using PP has previously been shown to induce calcium deposition in human aortic smooth muscle cells ^38^ and retinal pigmented epithelial cells ^39^, while organic phosphate using CM promotes cell mineralization and osteogenic differentiation of multipotent stem cells ^40^. The effects of this ectopically induced calcification by phosphate treatment, particularly in the context of mitochondrial calcium regulation, have yet to be evaluated by visualizing the structures of the RGC’s mitochondria under the pro-calcifying conditions.

In our study, we use cryoEELS and cryoET to directly characterize calcium granules in the mitochondria of RGCs treated with PP and CM. We establish a workflow for structurally evaluating and determining the elemental composition of in situ organelles in thin regions of intact cells at nanometer resolution. In addition, we used the same grid containing the vitrified RGC to visualize the 3D ultrastructure of mitochondria and applied a deep learning-based annotation and quantification system to estimate the size and distribution of the calcium granules. Our approach of correlating elemental mapping and 3D tomography of cells using cryoEELS and cryoET can be a powerful tool to explore disease phenotypes and mechanisms involving calcium depositions.

## Results

### Human embryonic stem cells differentiated into RGCs on electron microscopy grids

RGCs derived from hESCs offer a model system for studying mitochondrial calcification in its endogenous context. We optimized culture conditions for RGCs on Quantifoil® R 2/2 Micromachined Holey Carbon 200 mesh gold grids coated with poly-D-lysine and Laminin, achieving a proper cell density (5x10^5^ cells/ml) that allowed neurites to extend without crossing. The differentiation and identity of these RGCs were confirmed through quantitative PCR and tdTomato expression, which demonstrated the expression of Brn3B RGC markers (**Supplementary Figure 1**).

### CryoEELS reveals the presence of calcium in mitochondrial granules of RGCs treated with inorganic phosphate

A robust cryoEELS/cryoET workflow was established to investigate the intracellular elemental distribution and organelle structures in vitrified biological samples (**Figure 1**). We induced ectopic calcification in human RGCs using two different methods: one with inorganic phosphate (PP) and the other with organic phosphate (CM) (see Methods) ^37^. Given the mitochondria’s role in maintaining calcium homeostasis, we specifically targeted them in our analysis. Vitrified RGCs were examined in a 200kV JEOL electron microscope and screened to identify mitochondria in the thin neurites using Gatan’s Latitude S Software ^41^. After locating mitochondria that contained electron dense granules, a 2D cryoTEM reference image was taken, followed by STEM annular dark field (ADF) imaging and cryoEELS spectra acquisition.

**Figure 1.**
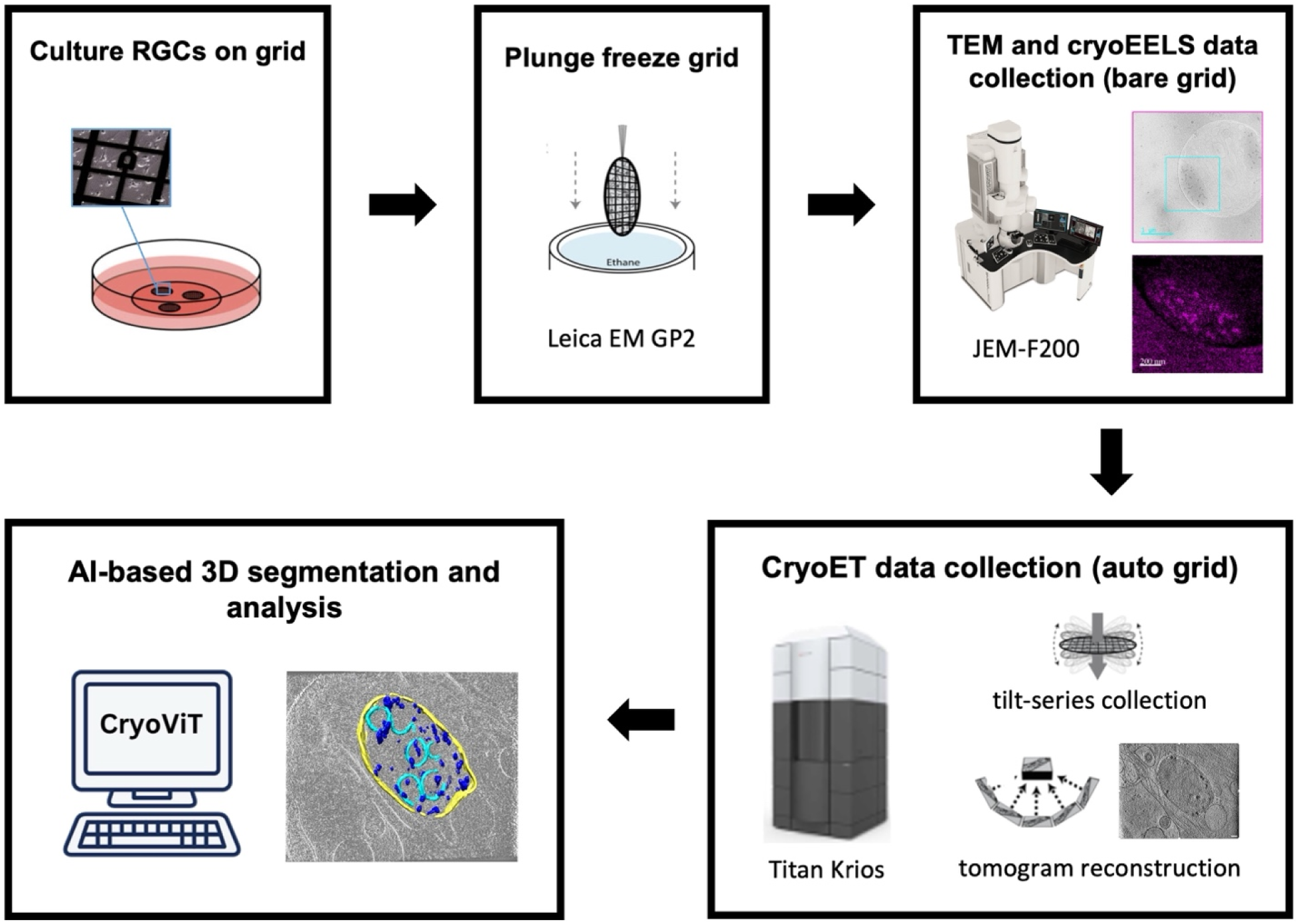
Workflow of cryoEELS and cryoET data collection. RGCs were cultured on the grid for 3 days, then plunge-frozen using a Leica GP2. 2D TEM images and cryoEELS data of the vitrified bare grid were acquired using a JEM-F200 equipped with a K3 camera. The same vitrified bare grid was then clipped to an autogrid and loaded into a ThermoFisher Titan Krios for cryoET data collection. Finally, an AI-based automatic segmentation tool was applied to the tomograms to visualize and quantify structural insights.

The carbon, calcium, nitrogen, and oxygen elemental maps reveal different elemental distributions in the subcellular structures in the vitrified RGCs treated with PP, as exemplified in two different regions found on the grid. The cryoTEM images of mitochondria reveal characteristic features of cristae as well as dark contrast granules (**Figures 2A, 2D**), whereas the ADF-STEM images display reverse contrast, with the granules appearing as bright spots (**Figures 2B, 2E**). The elemental maps for carbon, calcium, nitrogen and oxygen from these two regions are shown in **Figures 2C and 2F**.

**Figure 2.**
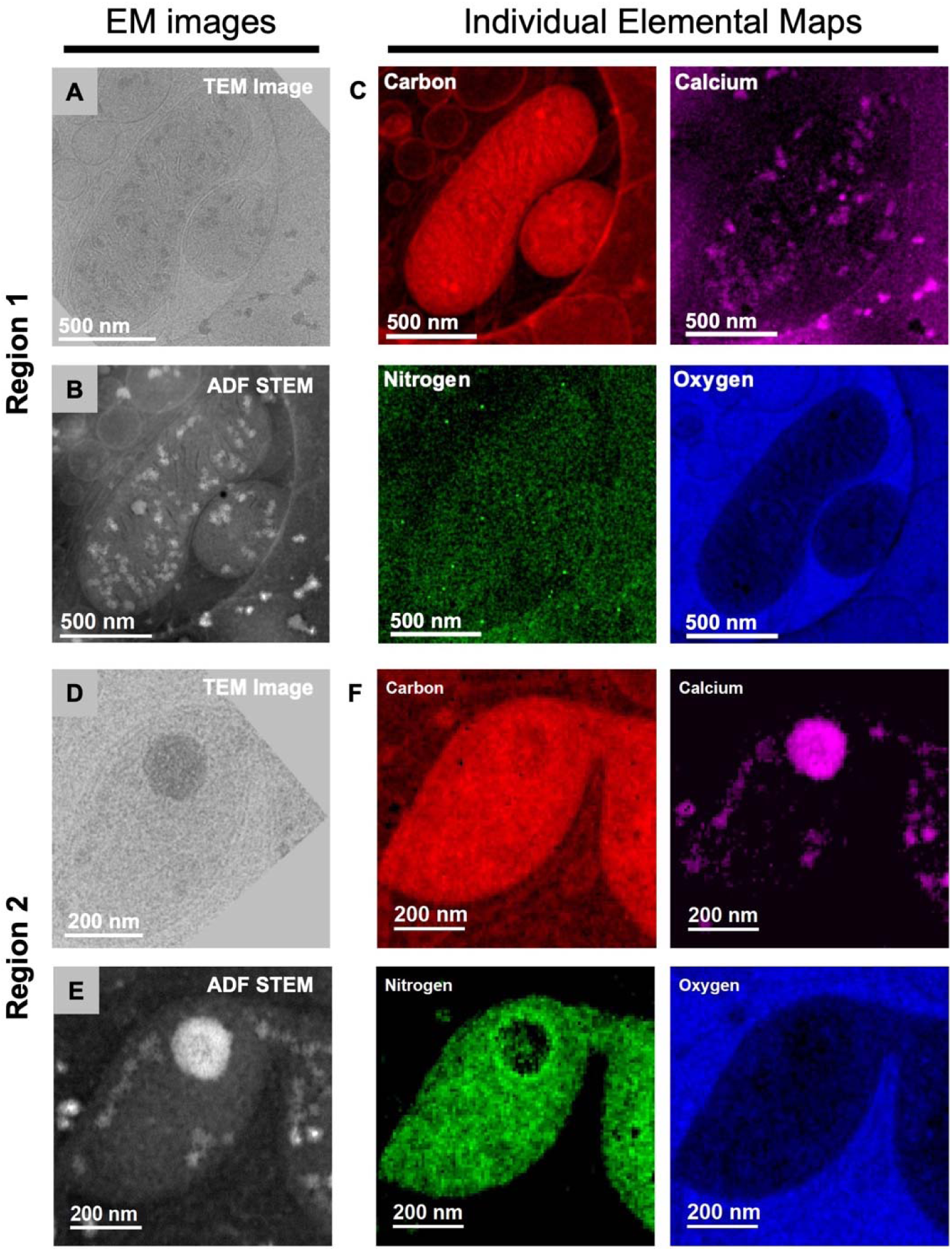
CryoEELS elemental analysis of two different mitochondria in potassium phosphate-treated retinal ganglion cells. (A) 2D TEM image reveals the double membrane, cristae, and granules of the mitochondria in region 1 of the grid. (B) Corresponding STEM-ADF image of the mitochondria in region 1 of the grid. (C) Individual elemental maps (carbon (K-edge), calcium (L_2,3_-edge), nitrogen (K-edge) and oxygen (K-edge)) of the mitochondria in region 1 of the grid. Scale bar is 500 nm. (D) 2D TEM image of the mitochondria with an enlarged granule in region 2 of the grid. (E) Corresponding STEM-ADF image of the mitochondria in region 2 of the grid. (F) Individual elemental maps (carbon (K-edge), calcium (L_2,3_-edge), nitrogen (K-edge) and oxygen (K-edge)) of the mitochondria in region 2 of the grid. Scale bar is 200 nm.

The carbon elemental maps (red contrast) are validated by their mapping to the carbon support film of the TEM grid, which can serve as a positive reference (**Figures 2C, 2F**). The high carbon signals (brighter red) could be contributed from the lipid membrane of cellular organelles such as vesicles, mitochondria, and mitochondrial cristae.

The brightest purple regions in the calcium elemental maps (**Figures 2C, 2F**) are consistent with the dark contrast features in the cryoTEM 2D image (**Figures 2A, 2D**) and the bright contrast features in the STEM-ADF image (**Figures 2B, 2E**) of the mitochondria containing electron dense granules. From the elemental maps in **Figures 2C and 2F**, calcium signals are also present outside the mitochondria. To further confirm the mapping between the dark field images and the elemental maps, we extracted the Ca spectra from different regions of the sample area as shown in **Figure 3**. Spectra extracted from the solid boxes in **Figures 3A and 3D** correspond to positive Ca L_2,3_ signals shown in **Figures 3B and 3E** (each line of the waterfall plot corresponds to the elemental signal from the matching colored box), respectively, while spectra extracted from the dashed boxes in **Figures 3A and 3D** correspond to negative Ca L_2,3_ signals shown in **Figures 3C and 3F**, respectively.

**Figure 3.**
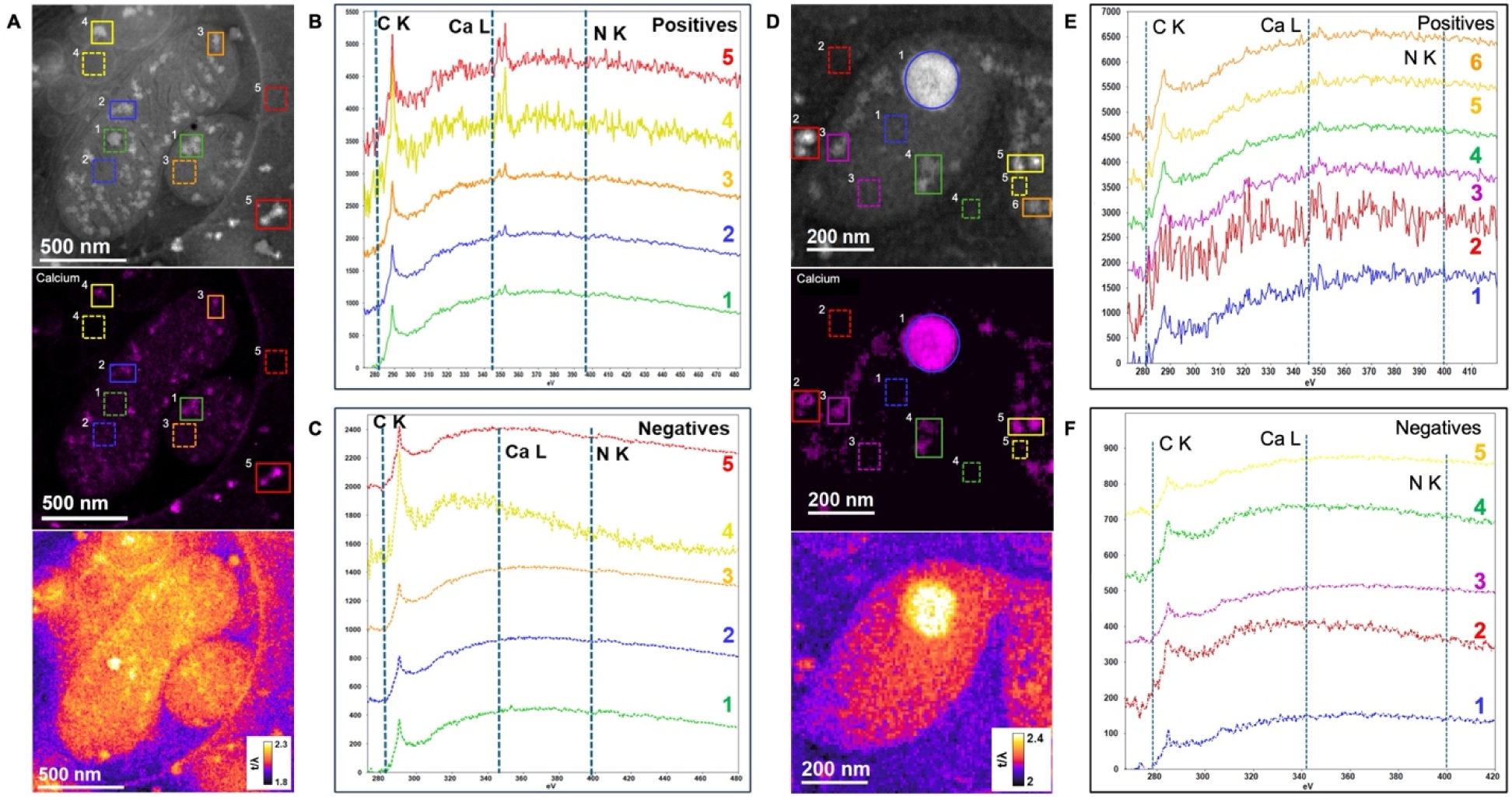
Calcium signal stack plot and inelastic mean free path image show calcium signals and sample thickness across mitochondria in retinal ganglion cells treated with potassium phosphate. (A) STEM-ADF image of the mitochondria in region 1 of the grid, where solid boxes indicate regions where the calcium L_2,3_ edge is detected (positive) and the dashed boxes indicate regions where the calcium L_2,3_ edge was not detected (negative). The middle of (A) shows the corresponding positive and negative regions in the calcium elemental map. At the bottom of (A) is a thickness map of the area showing how the sample thickness varies across the region, in units of inelastic mean free path (IMFP). (B) Waterfall plot of spectra extracted from positive regions in (A) where calcium was detected (solid boxes). (C) Waterfall plot of spectra extracted from negative regions in (A) where calcium was not detected (dashed boxes). (D) STEM-ADF image of the mitochondria in region 2 of the grid, where solid boxes indicate regions where the calcium L_2,3_ edge is detected (positive) and the dashed boxes indicate regions where the calcium L_2,3_ edge was not detected (negative). The middle of (D) shows the corresponding positive and negative regions in the calcium elemental map. At the bottom of (D) is a thickness map of the area showing how the sample thickness varies across the region, in units of inelastic mean free path (IMFP). (E) Waterfall plot of spectra extracted from positive regions in (D) where calcium was detected (solid boxes). (F) Waterfall plot of spectra extracted from negative regions in (D) where calcium was not detected (dashed boxes).

The nitrogen elemental maps (green contrast) vary in different mitochondria. A very weak nitrogen signal is observed in the mitochondria and the surrounding region in **Figure 2C**, whereas in **Figure 2F**, the nitrogen signal is localized to the mitochondria, with very little present in the surrounding region. The signal variation in the nitrogen elemental maps is likely due to variations in sample thickness. The N K-edge signal is observed to be relatively weak in the extracted spectra plotted in **Figures 3B, 3C, 3E, and 3F** and are barely detectable above the background. This is likely caused by the thick sample in these regions, which are calculated as shown in the bottom panel of **Figures 3A and 3D**. Notice that in the thickest regions of both samples, a relative thickness of 2.3 inelastic mean-free paths (IMFPs) is observed, meaning that each electron undergoes on average 2.3 inelastic scattering events as it travels through the sample. This increases the intensity of the extended edge region of the C K-edge which suppresses both the N K-edge signal and the Ca L_2,3_-edge signal.

The concentration of oxygen (blue contrast) does appear lower in the mitochondria than the surrounding regions in **Figures 2C and 2F**. This is due to the large amount of organic material in the mitochondria compared to the surrounding areas, which are composed mostly of vitreous ice. Therefore, in the mitochondria, inelastic scattering is more likely to occur from carbon compared to areas which are composed mostly of ice, where inelastic scattering is more likely to occur from oxygen.

From the elemental maps in **Figures 2C and 2F**, there appears to be no correlation between an increase in the Ca L_2,3_ edge intensity and an increase in the N K-edge or O K-edge intensities. However, the calcium granules may contain other biomolecules with nitrogen and oxygen atoms. In general, weak EELS signals can be detected by multiple least-squares modeling for the near-edge fine structure (ELNES) of an EELS ionization edge ^42^. However, due to the relatively weak N K-edge signal in the mitochondria, it would be impossible to perform multiple linear least squares fitting on the edge. Additionally, any potential O K-edge signal from the granules is likely suppressed by the large O K-edge signal originating from the surrounding ice.

### Mitochondria of RGCs cultured in organic phosphate-containing calcification medium (CM) have calcium granules

After establishing the cryoEELS workflow to detect calcium signals in mitochondrial granules of RGC neurites treated with PP, we applied this approach to analyze the elemental composition of RGCs treated with CM, which is typically used for osteogenic differentiation of mesenchymal cells. The carbon, calcium, nitrogen, and oxygen elemental maps revealed distributions comparable to those observed in RGCs treated with PP (**Figures 4, 5**). Notably, mitochondrial granules in CM-treated RGCs could also exhibit detectable calcium signals, indicating that CM treatment can induce mitochondrial calcification, which offers an alternative experimental protocol using CM treatment to study calcium homeostasis in mitochondria across various disease contexts.

**Figure 4.**
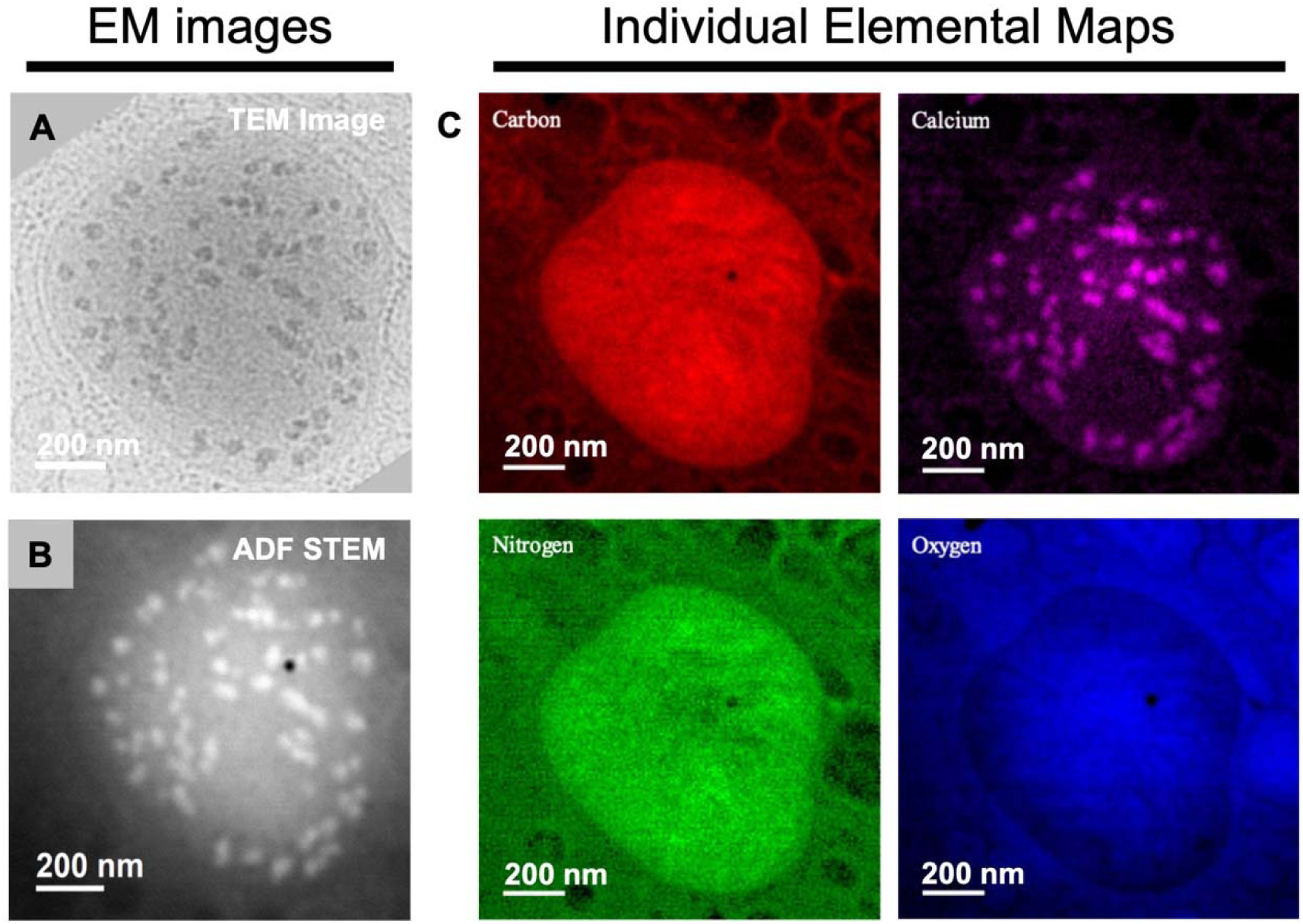
CryoEELS elemental analysis of a mitochondria in retinal ganglion cell treated with calcification media. (A) 2D TEM image reveals the double membrane, cristae, and granules of the mitochondria. (B) Corresponding STEM-ADF image of the mitochondria. (C) Individual elemental maps (carbon (K-edge), calcium (L_2,3_-edge), nitrogen (K-edge) and oxygen (K-edge)) of the mitochondria in region 1 of the grid. Scale bar is 200 nm.

There are some differences in the nitrogen and oxygen signals between the CM-treated and PP-treated RGCs, likely due to variation in sample thickness. In particular, the nitrogen signal (green) is relatively homogenous in **Figure 4C** and is observed in both the mitochondria, as well as the surrounding area. In contrast to the bottom panels of **Figures 3A and 3D**, the relative thickness of the region of the mitochondria in the bottom panel of **Figure 5A** is 1.5 IMFPs, and both the N K-edge signal and the Ca L_2,3_ edge signal have a higher signal-to-background ratio due to the reduction of plural scattering in a thinner specimen (**Figures 5B, 5C**). Additionally, in **Figure 4C**, the contrast in the oxygen map (blue) appears to be more homogenous relative to **Figures 2C and 2F**. It is likely that the mitochondria in this area are thinner, and that the top and bottom of that mitochondria are surrounded by a thicker layer of ice. This homogeneous distribution of contrast in the oxygen signal could also be due to the reduced suppression of the O K-edge caused by plural scattering.

**Figure 5.**
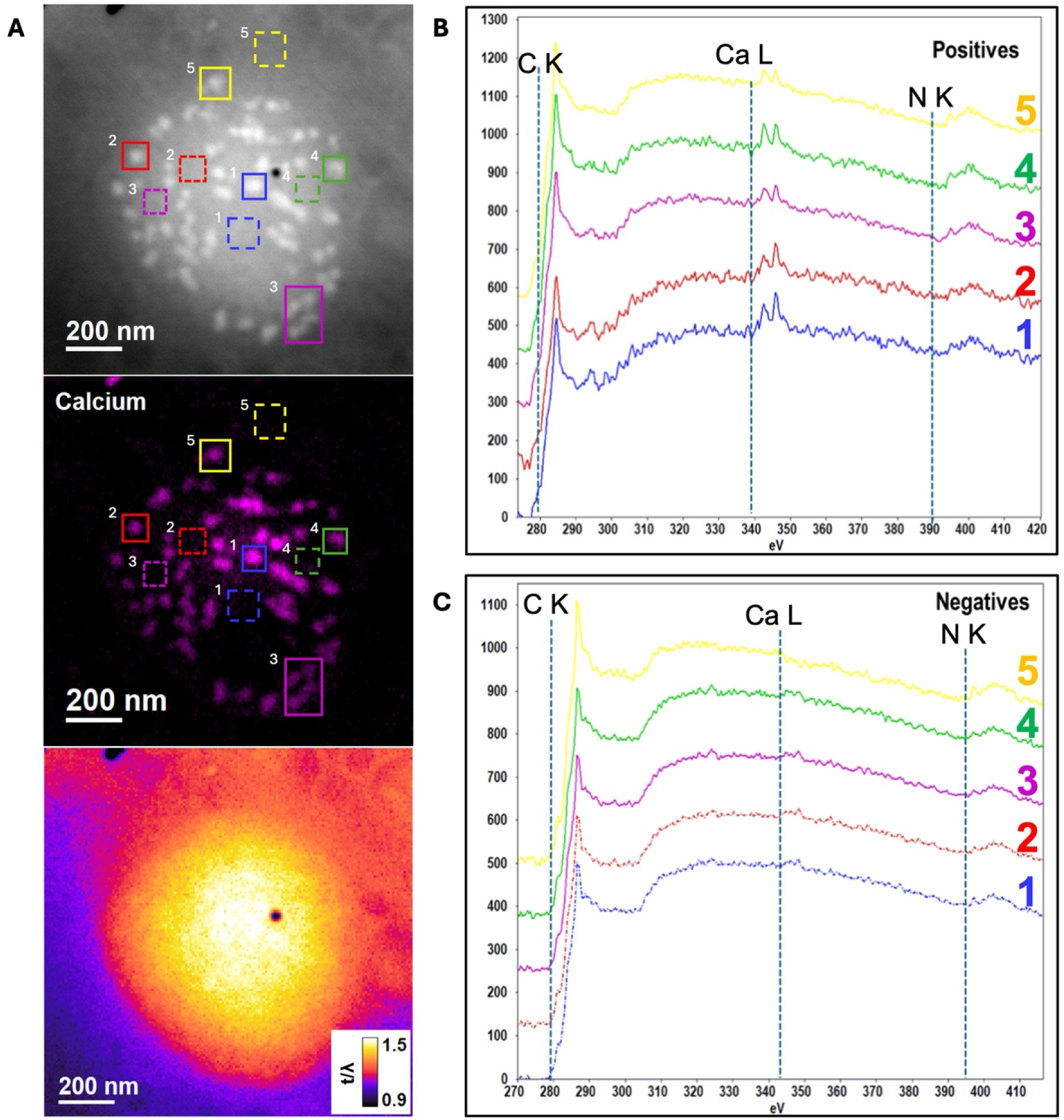
Calcium signal stack plot and inelastic mean free path image shows calcium signals and sample thickness across mitochondria in retinal ganglion cells treated with calcification media. (A) STEM-ADF image of the mitochondria, where solid boxes indicate regions where the calcium L_2,3_ edge is detected (positive) and the dashed boxes indicate regions where the calcium L_2,3_ edge was not detected (negative). The middle of (A) shows the corresponding positive and negative regions in the calcium elemental map. At the bottom of (A) is a thickness map of the area showing how the sample thickness varies across the region, in units of inelastic mean free path (IMFP). (B) Waterfall plot of spectra extracted from positive regions in (A) where calcium was detected (positive). (C) Waterfall plot of spectra extracted from negative regions in (A) where calcium was not detected (negative).

### Artificial intelligence-based segmentation enabled quantification of mitochondrial granules

To further investigate the effects of PP and CM treatment on RGCs, we collected and reconstructed cryoET tomograms from the same grid that were used for the cryoEELS data collection. In total, we obtained 26 tomograms from PP-treated RGCs and 36 tomograms from CM-treated RGCs, with examples shown in **Figures 6A and 6B**. Electron-dense granules were observed in the mitochondria of RGCs under both the PP and CM treatment conditions. To analyze the volume and density of these mitochondrial granules, we developed a deep-learning-based segmentation and quantification method. We segmented a total of 90 mitochondria and 1295 mitochondrial granules from 11 tomograms of PP-treated RGCs (**Supplementary Movie 1**) and 27 tomograms of CM-treated RGCs (**Supplementary Movie 2**).

**Figure 6.**
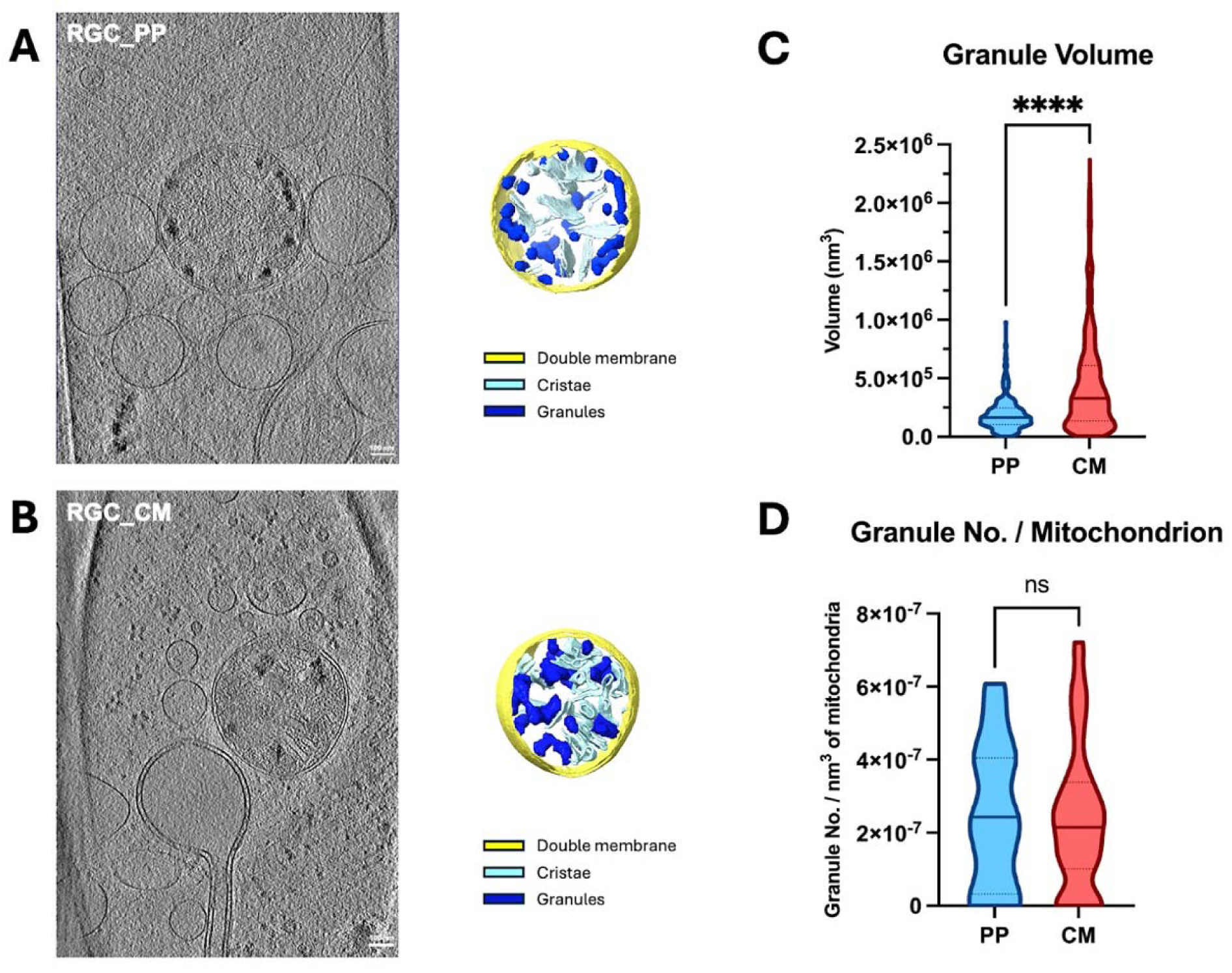
AI-based mitochondrial granules annotation reveals the enlarged calcium granules in retinal ganglion cells treated with potassium phosphate and calcification media. (A) Single slice of cryoET image reveals the mitochondrial morphology and mitochondrial granules in PP-treated RGC mitochondria. Semi-automated annotation reveals the double membrane, cristae, and granules of the mitochondria. (B) Single slice of cryoET image reveals the mitochondrial morphology and mitochondrial granules in CM-treated RGC mitochondria. Semi-automated annotations reveal the double membrane, cristae and granules of the mitochondria. (C) Mitochondrial calcium granules in CM-treated RGCs are larger than those in PP-treated RGCs. (D) Mitochondrial granule density is the same between CM- and PP-treated RGCs. ****=p < 0.0001. ns=no significant difference.

Quantification of mitochondrial granule volumes using our segmentation method ^43^ revealed that the size distribution of granules in CM-treated RGCs was shifted towards larger sizes compared to PP-treated RGCs (**Figure 6C**). Although the average granule volume increased with CM treatment, the number of granules per nm³ of mitochondrial volume remained similar between the two conditions (**Figure 6D**). In PP-treated RGCs, the average granule volume was 1.94x10^5^ ± 1.40x10^5^ nm³, and the average number of granules per nm³ of mitochondria was 2.46x10^-7^ ± 1.99x10^-7^. For CM-treated RGCs, the average granule volume was 4.50x10^5^ ± 4.21x10^5^ nm³, and the average number of granules per nm³ of mitochondria was 2.40x10^-7^ ± 1.97x10^-7^. These findings suggest that both PP and CM treatment of RGCs induces the formation of calcium granules in mitochondria and can serve as *in vitro* models to study mitochondrial calcification in the context of ODD.

## Discussion

In this study, we have directly characterized the composition of mitochondrial granules in situ using cryoEELS and cryoTEM, achieving unprecedented spatial resolution. Our approach involves efficiently and rapidly scanning the grid in cryoTEM mode on a STEM instrument using Gatan’s Latitude S software to identify areas containing mitochondria with double membranes, cristae, and granules ^41^. This is followed by simultaneous ADF and cryoEELS spectra acquisition to generate elemental maps of carbon, calcium, nitrogen, and oxygen. The regions of the elemental maps corresponding to visible granules in the TEM and STEM-ADF images exhibited high calcium signals, confirming that these granules are indeed composed of calcium. This direct observation of calcium granules within mitochondria in their natural environment provides insight into how calcium is stored in these organelles. It also suggests that dysregulation of calcium levels and an overload of calcium granules may contribute to dysfunctional mitochondria and lead to pathogenesis. Furthermore, the elemental maps revealed intricate details of other subcellular structures. For example, the carbon elemental map clearly delineated the outline of carbon-rich lipid membranes, including the mitochondrial membrane, cristae, and single-membrane vesicles. The lower oxygen concentration in the mitochondrial matrix relative to other areas is attributable to the fact that most of the sample thickness in the mitochondrial area is made up of cellular material (carbon) rather than ice, which contributes substantially to the oxygen signal.

Sample thickness is a crucial factor in cryoEELS analysis. As sample thickness increases, the electrons interact more with the sample, which results in multiple scattering effects ^44^. This phenomenon tends to reduce the signal-to-noise ratio of EELS edges, diminishing their detectability. Initially, we intended to work with fibroblasts, which were more readily accessible. However, the fibroblast thickness was estimated to be 400-500 nm by cryoET, which resulted in very weak elemental signals in our cryoEELS experiment. To overcome this challenge, we used hESC-derived RGCs, which contain neurites that are sufficiently thin (200-300 nm) for high-signal-to-noise ratio data collection of the cryoEELS SI in addition to its relevance to ODD. To work with thicker specimens, such as whole cells or tissues, while maintaining molecular resolution, we would need to employ cryogenic focus ion beam (FIB)-milling ^45^. This method allows for the preparation of thin lamellae with a final thickness of 100-200 nm from thicker samples, making them suitable for high-resolution cryoET and cryoEELS.

Structural analysis of the mitochondrial granule distribution revealed a similar number of granules in RGCs treated with CM compared to those treated with PP, but the granules in CM-treated RGCs were larger on average. High inorganic phosphate may induce calcium phosphate deposits due to its propensity to complex with cations like calcium and magnesium or bind to proteins ^46^. The CM cocktail, typically used for osteogenic differentiation of mesenchymal stem cells, consists of dexamethasone (Dex), ascorbic acid (Asc), and β-glycerophosphate (β-gly). The transcriptional activation in cells influenced by CM treatment affects intracellular signaling and promotes calcium phosphate crystallization to form hydroxyapatite ^40^. Specifically, Dex induces the expression of Runx2, a potent calcification inducer, through the transcriptional activation of FHL2/β-catenin and the upregulation of TAZ and MKP1. Asc increases the secretion of collagen type I (Col1), which enhances Col1/α2β1 integrin-mediated intracellular signaling. β-Gly serves as an organic phosphate source for hydroxyapatite and influences intracellular signaling molecules. The larger mitochondrial granules observed in CM-treated RGCs suggest that the transcriptional activation induced by the CM cocktail promotes gradual and greater calcification compared to extreme and direct inorganic phosphate stress in the PP condition. Phosphate in both treatments resulted in mitochondrial granules composed of calcium, indicating that PP- and CM-treatment can serve as models to study RGC mitochondrial calcification for ODD. These findings also highlight that cryoET structural quantification alone is insufficient to determine the chemical composition of the targeted subcellular ultrastructure, emphasizing the importance of combining measurements from both cryoET and cryoEELS.

There is a clear advantage to performing both ultrastructural and elemental characterization of the same grid area within a single electron microscope, provided it is equipped with the necessary energy filters and detectors for both cryoEELS and cryoET experiments. Unfortunately, cryoEELS is generally configured for STEM in electron microscopes optimized for materials science applications, whereas cellular cryoET is performed in microscopes optimized for biological specimens. In our experiment, we successfully performed cryoEELS and cryoET on the same grid using two different electron microscopes. However, we were unable to use the same region of interest for both measurements due to two main factors: the difficulty in relocating specific regions across a series of experiments and the relatively high electron dose used in cryoEELS, which damages the sample and renders it unusable for subsequent cryoET data collection.

Ideally, we would capture cryoET images in the Titan Krios and then perform cryoEELS analysis in another STEM instrument. However, this approach is currently hindered by the incompatibility between the AutoGrids used in the Titan Krios and the cryo-holders in side-entry STEM instruments. This incompatibility significantly increases the risk of disrupting the bare grid during transfer, making it difficult to collect both cryoEELS and cryoET data from the same grid, which is essential for maintaining sample consistency and obtaining measurements from the same cellular targets. Moreover, performing cryoET before cryoEELS is currently impossible due to the inability to remove the vitrified grid from the AutoGrid cartridges without causing damage.

In order to address these technical issues and optimize our workflow, it is essential to develop a side-entry holder compatible with AutoGrids. This development would enable cryoET data collection before cryoEELS spectra acquisition from the same grid area. Such an approach offers several advantages, including the ability to use low-dose search and imaging modes with cryoTEM to locate targets of interest in vitrified cells prior to the final cryoEELS step, which typically requires a higher electron dose. Additionally, it would allow the protocol to be extended to integrate prior experiments using cryoFIB-SEM and cryoCLEM on vitrified thick cells or entire tissues before following the workflow established in this study.

Furthermore, this integration of structural and compositional analyses at the subcellular level can be adapted to study the presence of various elements in different organelles. In our previous study, we observed enlarged mitochondrial granules in iPSC-derived neurons from Huntington’s disease (HD) patients and HD mouse primary neurons ^47^. Proteomics analysis revealed increased RNA binding proteins, leading us to hypothesize that those granules were mitochondrial RNA granules (MRGs). Knockout (KO) of GRSF1, an RNA binding protein found in MRG aggregates, resulted in reduced granule sizes. However, both healthy samples and GRSF1 KO samples still exhibited some remaining small granules, which can be hypothesized as calcium phosphate granules. Further studies utilizing our correlated cryoEELS and cryoET workflow could further elucidate the composition and nature of these residual granules. Identifying abnormal subcellular structures and their compositions provides valuable insight into the underlying mechanisms of their dysregulation. For example, this workflow could be applied to investigate iron accumulation in neurodegenerative disorders such as Alzheimer’s disease ^48^. By combining high-resolution imaging with elemental analysis, we can gain unprecedented insights into cellular landscapes in both healthy and disease states. This multimodal approach not only advances our understanding of cellular biology but also holds promise for identifying novel therapeutic targets and biomarkers across a wide range of pathological conditions.

## Methods

### Ethical statement

All experimental work performed in this study was approved by the Human Research Ethics Committee of Stanford University (SCRO#0875).

### Human embryonic stem cells maintenance and differentiation to RGCs

Human embryonic stem cells (hESCs) with the BRN3B–P2A-tdTomato-P2A-THY1.2 construct gifted by Professor Donald Zack (Johns Hopkins University) were used in our study. These hESCs were maintained as described ^49^. Briefly, cells were cultured on growth factor-reduced Matrigel-coated plates in mTeSR1 media at 5% CO_2_. hESC colonies were passaged using Accutase or gentle cell dissociation reagent for 3 minutes at 70-80% confluency as small clumps, split in a 1:4 ratio every 5-7 days, and maintained at 37°C in a 5% CO_2_ incubator. hESC differentiation to retinal ganglion cells (RGCs) was obtained by using small molecules as previously described ^49^. Single cells were generated using Accutase for 10 minutes and cultured in mTeSR1 media with 10 μM ROCK inhibitor on Matrigel-coated plates (defined as day -1). The following day, the medium was completely changed to RGC differentiation basal media 1 (RDBM1), comprising DMEM/F12 and Neurobasal (1:1) with GlutaMAX, 1X penicillin-streptomycin, 1% N2 Supplement, and 2% B27 Supplement (defined as day 0). From day 1 to day 6, the following small molecules were added to RDBM1: Forskolin (FSK; 25 μM), Dorsomorphin (DSM; 1 μM), IDE2 (2.5 μM), and Nicotinamide (NIC; 10 mM), with daily media changes. From day 6 to day 10, cells were treated with FSK and NIC only; from day 10 to day 18 with FSK only; from day 18 to day 30 with FSK and DAPT (10 μM); and from day 40 onwards with RDBM1 only. Successful RGC differentiation was monitored microscopically for tdTomato expression.

### Purification and magnetic sorting of RGCs

RGCs were purified between day 45 to day 50 by dissociating differentiated cells with Accutase, followed by sorting using CD90.2 (THY1.2) microbeads and a magnetic activated cell sorting system (MACS, Miltenyi Biotec) as described by Sluch et al. ^49^. CD90.2 microbeads were added to the cell suspension according to manufacturer instructions and incubated for 15 minutes at 4°C for cell binding. To increase RGC purity, cells were first passed through an LS column. The flow-through comprising negative cells was discarded, while positive cells attached to the column were flushed out of the media with a plunger. These positive cells were then passed through an MS column. Following sorting, the cells were maintained in N2B27 medium supplemented with 10 mM FSK and 10 ng/ml CNTF on Matrigel-coated plates and were used for various experimental conditions. Sorted RGCs were tdTomato positive and were further characterized by qPCR for RGC markers (Brn3b/Gapdh).

### Cell growth conditions and vitrified grid preparation

Quantifoil® R 2/2 Micromachined Holey Carbon grid: 200 mesh gold (SPI supplies; Catalog No. 4420G-XA) grids were treated with 0.1 mg/ml poly-D lysine (Fisher Scientific; Catalog No. A3890401) at 4°C overnight in a 35 mm Mat-Tek glass coverslip bottom dish (VWR; Catalog No. P35G-0-14-C). After the poly-D lysine coating, the grids were washed three times with double-distilled water, air dried, and further coated with Laminin (3ug/mL). 1x10^5^ magnetic sorted RGCs were cultured on the carbon side of the coated grids for 3 days in N2B27 medium supplemented with 10 mM FSK and 10 ng/ml CNTF.

Cells cultured on the grids were vitrified using the temperature- and humidity-controlled Leica GP2 (Leica Microsystems). The grids were loaded and blotted for 7 seconds at 37°C and 95% humidity, then immediately plunged into liquid ethane. The vitrified grids were subsequently transferred to storage boxes containing clean liquid nitrogen, where they remained until cryoET data collection.

### CryoEELS spectra data collection

CryoEELS spectrum images (SI) were acquired on a JEOL F200 (Jeol USA) equipped with a Cold Field Emission Source in scanning TEM (STEM) mode. The suppressor and extractor were set to 3.85 keV and 7 keV, respectively, to achieve an emission current of 10 μA. A Gatan 910 cryo-transfer holder was used to maintain a sample temperature of 170°C during the loading of the sample into the microscope and during image and cryoEELS acquisition. Due to the thickness of the sample and the small difference in atomic weight between oxygen, nitrogen, and carbon, the STEM annular dark field (ADF) images have very low contrast. Searching for specific mitochondria of interest was extremely difficult due to the low contrast. Therefore, a low-magnification bright field TEM image was used to identify key regions of interest. Then, the microscope was switched into STEM mode. STEM ADF images and cryoEELS SI were acquired simultaneously at spot size #9, producing a probe current of 3.7 pA. A defocus of -5 nm was used to set the STEM probe diameter to 1 nm. CryoEELS SI were acquired with a convergence semi-angle (α) of 10 mrad and a collection semi-angle (β) of 31 mrad on Gatan’s Continuum HR spectrometer (Gatan) with a K3 direct detection camera ^29^, with a dispersion of 0.45 eV/ch. DualEELS was used to incorporate plural scattering into the spectral models to ensure the most accurate fit. Multipass spectrum imaging was used to fractionate the total dose over multiple passes to reduce the effects of beam damage^50^. Multipass SI was acquired using the DigitalMicrograph in-situ spectrum imaging tool (Gatan), which allowed each pass to be saved individually rather than as a continuous sum. Acquiring individual passes had two key advantages. First, due to the small camera length and low probe current, the image intensity on the ADF image prohibited automated drift correction. Since frames were saved individually, drift was corrected manually post-acquisition. Secondly, mitochondria are significantly dose sensitive. Saving individual passes allowed the removal of SI passes in which mitochondria were compromised by radiolysis. CryoEELS SI were acquired with a pixel size of 8 nm and a pixel dwell time of 4 ms. Sub-pixel scanning was used to spread the dose over the entire pixel area, using an 8x8 subpixel array to match the 1 nm probe diameter. Although subpixel scanning does not increase cryoEELS SI resolution, it reduces the total dose per subpixel area. In this experiment, the dwell time per subpixel was 62.2 μsec. Together with a probe current of 3.7 pA, these conditions gave a total dose per SI pass of 18.4 e^-^/Å^2^.

### CryoEELS analysis of calcium (Ca) in mitochondrial granules

After acquisition, cryoEELS SI were spatially aligned using the DigitalMicrograph In-situ tools (Gatan) to correct for spatial drift that occurred during measurement. A Sobel filter was used to enhance edges in the SI to measure drift between passes. After spatially aligning each pass, the stack of simultaneously acquired ADF images was scanned to detect void formation (bubbles) in the specimen ^27^. Passes up to, but not including the point of void formation, were kept and summed to form the final SI, while remaining passes were discarded. It was found that void formation occurred at an accumulated dose of about 315 e^-^/Å^2^, so a total of 15 spectrum image passes were summed to form the final SI. CryoEELS elemental maps were calculated using model-based quantification ^51^. The model comprises a linear combination of components, with multiple linear least squares fitting used to obtain the best coefficient of fit. A standard power-law was used to model the background under the ionization edges. Two separate models were used to model the ionization edges ^14^. The Hartree-Slater cross-sections were used to model the extended edge fine structure of each edge ^52^. Since the Hartee-Slater cross sections do not consider solid-state effects, the near-edge fine structure (ELNES) was modeled using internal and external references. The ELNES model for the C K-edge was extracted from an amorphous carbon support film (Ted Pella Inc), and the Ca L_2,3_ edge model was extracted from CaCO_3_ particles supported on a 25-nm thick SiN membrane. These references were acquired at 0.45 eV/ch dispersion, with plural scattering removed using the Fourier-log method ^14^. Since the shape of the O K-edge originates from vitrified ice, an internal ELNES reference was acquired from a region where the ice spans a hole in the Quantifoil® carbon film. Samples were calculated to be 150-200 nm thick, and a significant contribution to the edge intensity was due to plural scattering. The model was convoluted with the low-loss portion of the EELS spectrum to account for plural scattering. This was especially critical for extracting the Ca L_2,3_ signal since plural scattering from the extended edge region of the C K-edge significantly increases the background under the Ca edge. It is also worth noting that above a sample thickness of 200 nm, the plural scattering from the C K-edge likely quenches the Ca L_2,3_. Although we could detect similar bright areas in the ADF image, which corresponded to similar contrast observed in calcium granules observed in thinner areas, calcium was undetectable in samples above this thickness.

### CryoET data collection, tomographic reconstruction, and annotation

We used cryoET to characterize the RGCs with PP and CM treatment from the same grids after collecting the cryoELLS. Tilt series were collected in a Titan Krios electron microscope (ThermoFisher) equipped with a BioQuantum Energy Filter (set to 20 eV) and operated at 300 kV in low-dose mode using SerialEM software ^53^. At each tilt angle, we recorded “movies” with 5 frames each using a K3 camera (Gatan). The tilt series were collected dose-symmetrically from 0° to ±60° with a 2° tilt step, target defocus of -5 μm, and a cumulative dose of ∼121 e/Å^2^. Images were motion-corrected with MotionCor2 software ^54^. We used the IMOD software package ^55^ for standard weighted-back projection tomographic reconstruction after patch-tracking alignment. Within each tilt series, unusable images (due to large drift, excessive ice contamination, etc.) were removed prior to alignment. Data were corrected with a contrast transfer function (CTF) using IMOD’s 3D-CTF correction algorithm ^56^. Reconstructed tomograms were binned by 4 and post-processed using low-pass and high-pass filters, normalization, and thresholding at 3 standard deviations from the mean. Tomographic annotation of membranes and other features in binned-by-4 tomograms was carried out using EMAN2’s semi-automated 2D neural network-based pipeline ^57^ and MemBrain-seg ^58^. Manual clean-up of false positives was performed in ChimeraX ^59^, which was also used to display all annotated densities and prepare cryoET movies for structural interpretations.

### Quantification and statistical analysis of mitochondrial granule distribution

We employed a two-stage deep learning system for automatic voxel-level annotation of mitochondria and granules in tomograms ^43^. In the first stage, our system is trained on a small subset of annotated slices—only 2% of the 2D slices from each tomogram were manually annotated, with each pixel labeled as either background or part of a mitochondria or granule. In the second stage, we used our trained model to generate predictions to fill in the remaining unannotated slices in each tomogram. We trained two separate models using this approach: one for segmenting mitochondria and another for segmenting granules. We kept the granule annotations that were located within a detected mitochondrion and manually cleaned up false positives using ChimeraX ^59^. Using Segger ^60^, we identified individual instances of granules and mitochondria from the predictions and quantified their numbers and volumes within the tomogram. We scaled each voxel accordingly to obtain the volume in nm^3^.

Data was compiled into graphs using GraphicPad Prism 10. The data are presented as mean ± standard deviation. Only tomograms containing mitochondria with electron-dense granules were included for the analysis. The Mann–Whitney U-test was used to assess differences between the PP and CM treatment groups. A p-value < 0.05 was assumed to be statistically significant.

## Acknowledgements

This research has been supported partially by Stanford Wu Tsai Neurosciences Institute (KIG-102).

## Author Contributions

This project was conceived by W.C. with advice from Y.J.L. on the choice of benchmark samples and technical guidance for cryoTEM and cryoEELS from S.G. and C.B. Sample preparation was performed by H.R.P. and G.-H.W. The cryoEELS experiments and analyses were carried out by A.T., C.B. and G.-H.W. The cryoET experiments were carried out by G.-H.W. and C.H. Tomographic reconstructions, annotations, and statistical analyses were done by C.H. with feedback from G.-H.W. and S.R.G. The figures and movies were prepared by C.H. with feedback from G.-H.W., A.T. and W.C. The manuscript was written by C.H., A.T., G.-H.W., H.R.P. and W.C. with major input from Y.J.L. and feedback from all authors.

## Data Accessibility

Representative tomograms for each condition investigated in this study have been deposited in the EMDB (https://www.emdataresource.org/deposit.html). The accession codes are as follows: EMD-XXX (human retinal ganglion cell treated with potassium phosphate) and EMD-XXX (human retinal ganglion cell treated with calcification media).

---

**Figure S1.**
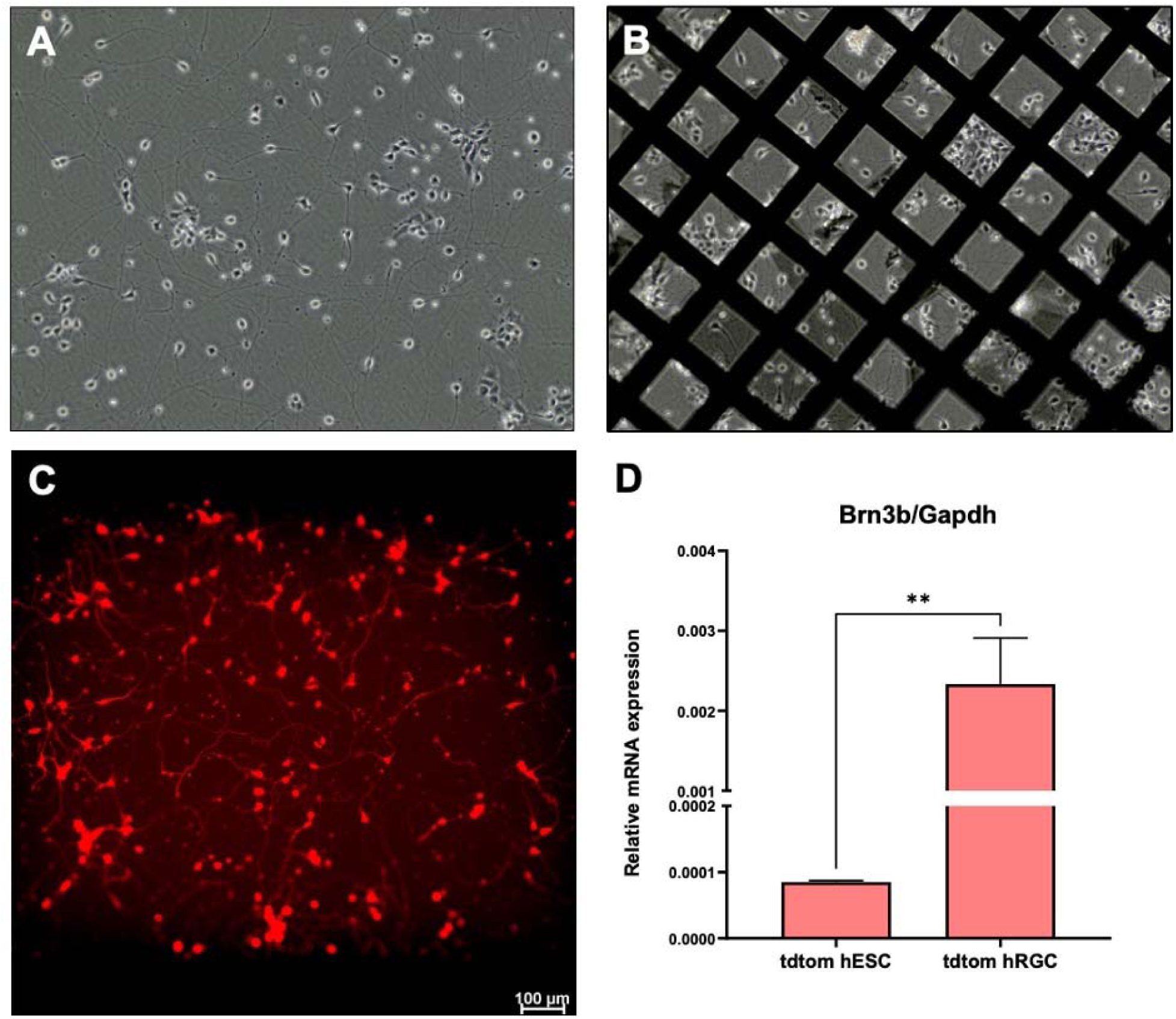
TdTomato+ human retinal ganglion cells grid culture conditions were monitored and differentiation was identified. (A) Bright field image of tdtom hRGCs growing on the glass. (B) Bright field image of tdtom hRGCs growing on Quantifoil® R 2/2 Micromachined Holey Carbon 200 mesh gold grid. (C) Fluorescent image of tdtom hRGCs growing on the glass. (D) Increased RNA expression of RGC marker *brn3b* in induced tdtom hRGCs compared with human embryonic stem cells.

